# Multi-view BLUP: a promising solution for post-omics data integrative prediction

**DOI:** 10.1101/2024.11.14.623556

**Authors:** Bingjie Wu, Huijuan Xiong, Lin Zhuo, Yingjie Xiao, Jianbing Yan, Wenyu Yang

## Abstract

Phenotype prediction is a promising strategy for accelerating molecular plant breeding. Data from multiple sources (called multi-view data) can provide complementary information to characterize a biological object from various aspects. By integrating multi-view information into phenotype prediction, a multi-view best linear unbiased prediction (MVBLUP) method was proposed in this paper. By assigning different weights to measure the importance of different views of data and using a differential evolution algorithm with an early stopping mechanism to adjust the weight vector adaptively, a multi-view kinship matrix was obtained first and then incorporated into the BLUP model for phenotype prediction. To validate the efficiency of MVBLUP, we conducted numerical experiments on four multi-view datasets.

Compared to the average performance of the single-view method, the prediction accuracy of the MVBLUP method has improved by a range of 0.038 to 0.201.The results demonstrate that the MVBLUP model is an effective method of integrating multi-view data.

## Introduction

Phenotype prediction is a powerful tool that allows for the early assessment of traits in individuals prior to planting, thereby accelerating the breeding process and significantly reducing its duration. This prediction is primarily achieved through genomic prediction (GP), a concept initially introduced by Meuwissen for animal breeding (Meuwissen et al., 2001). Since then, various methods have been employed to predict traits, broadly categorized into traditional statistical methods and machine learning approaches. Traditional statistical methods encompass best linear unbiased predictions (BLUP) (Henderson, 1975), least absolute shrinkage and selection operator (LASSO) (Usai et al., 2009), and Bayesian-based methods such as Bayes A, Bayes B, Bayesian LASSO (Meuwissen et al., 2001; Yi and Xu, 2008; de los Campos et al., 2009). On the other hand, machine learning approaches include support vector machine (SVM) (Maenhout et al., 2007), random forest (RF) (Holliday et al., 2012), deep convolutional neural networks (DeepGS) (Ma et al., 2018), and deep neural network for genomic prediction using multi-omics data (DNNGP) (Wang et al., 2023). Notably, DNNGP incorporates a batch normalization (BN) layer to mitigate overfitting and can be viewed as an advanced version of DeepGS. Other notable approaches include extreme gradient boosting (XGBoost) (Chen and Guestrin, 2016; Xu et al., 2016) and its faster version, light gradient boosting machine (LightGBM) (Yan et al., 2021).

Multi-view data refers to data from multiple sources that offer complementary information to characterize a biological object from various perspectives. These data can include different groups of samples measured by the same feature set (multi-class data), the same samples with various feature sets (e.g. multi-omics data), the same samples by the same set of features under different conditions (e.g., multi-environmental data), or different features and different samples in the same system (multi-relational data) (Li et al., 2018). Such data are ubiquitous in the real world. For example, a sample can be characterized by its genotype, gene expression levels and metabolomic data, each serving as a unique view of the sample. Compared to single-view data, multi-view data can provide more complementary information, and thus, effective integration of multi-view data has the potential to enhance model prediction performance (Serra et al., 2015; Dimitrakopoulos et al., 2017; Montesinos-López et al., 2018).

Recent studies have demonstrated that integrating multi-view data can lead to higher prediction accuracy. For example, incorporating transcript levels from seedlings with genetic markers into a joint model improved the prediction of mature maize traits (Azodi et al., 2019). By integrating metabolomics data into genomic prediction of hybrid yield of rice, predictability was enhanced by approximately 30% (Xu et al., 2016). During the learning process of a LASSO model with genome-wide markers, sequential integration of transcriptome and metabolome features allowed for iterative learning of three layers of features, resulting in significant improvement in rice yield trait prediction (Hu et al., 2019). Additionally, incorporating genotype×environment interaction (G×E) into a GP model has been shown to improve prediction accuracy (Montesinos-López et al., 2018; Crossa et al., 2021; Xu et al., 2022; Barreto et al., 2024).

Despite the demonstrated efficiency of multi-view integration prediction, there is still considerable room for improvement in learning from different data views. The relationship among different views is often complex, with different data sources potentially containing varying amounts of information and noise. The quality of data typically varies across different samples, meaning that one view may be informative for one sample but not for another. Existing multi-view methods often treat each view with equal importance, tune their weights to fixed values, or integrate them with a black-box machine learning framework (Wang et al., 2021). Therefore, there is a need to develop new methods for integrating multi-view information.

The differential evolution algorithm (DE), first proposed by Storn and Price, is a population-based evolutionary algorithm (EA) designed to search for a parameter set that maximizes a target function (called as fitness function) (Storn and Price, 1997). The algorithm mimics natural evolution through an iteration process involving mutation, crossover and selection, evolving the population towards better solutions. In the mutation phase, DE generates a new individual (called the mutant vector) by computing the difference between two randomly selected individuals, scaling the difference by a factor and adding the result to a third randomly selected individual. At the crossover stage, a trial individual can be generated by combining the mutant individual with a target individual with a certain probability determining which genes come from the mutant and which from the target individual. The selection step evaluates the fitness value of the trial individual and compares this fitness value with that of the target individual. If the fitness of the trial individual is better, replace the target individual with the trial individual in the population. Otherwise, keep the target individual. It ensures that only individuals with improved fitness are retained in the population.

In this study, we adapted DE to establish an adaptive multi-view integration strategy to better measure the importance of different views. By combining this adaptive integration strategy with the common statistical prediction model, BLUP, we proposed a multi-view best linear unbiased prediction (MVBLUP) method for phenotype prediction. The schematic workflow of the method is illustrated in Fig. 1, and details of the algorithm can be found in the Method subsection of the paper. To evaluate the performance of the MVBLUP, we compared its prediction accuracy, with BLUP, LASSO and XGBoost using single-view and multi-view data from tomato, rice and maize datasets of different sizes. Numerical results demonstrate that MVBLUP is a promising and practical approach for integrating multi-view data for phenotype prediction.

**Fig. 1.**
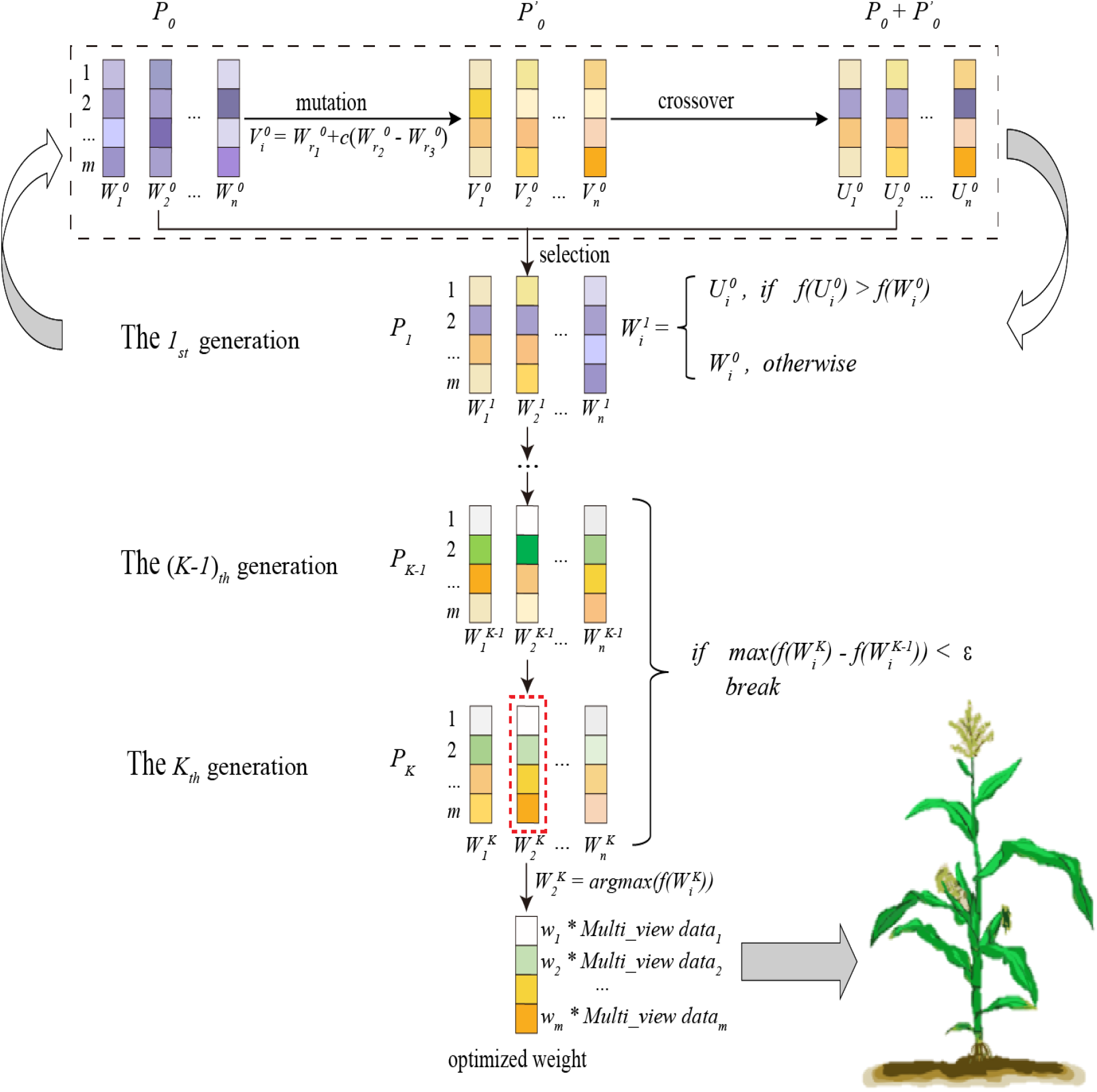
The schematic workflow of the MVBLUP algorithm, which entails the adaptive learning of optimal weights reflecting the significance of each view via DE algorithm. These learned weights are then employed to construct a multi-view kinship matrix for the MVBLUP model.

## Results

A comparative analysis of the results was conducted by using MVBLUP, BLUP, LASSO and XGBoost, each evaluated separately on the datasets Tomato332, Rice210, Maize368, and Maize282. The standard for assessing these comparison results was the average prediction accuracy derived from 50 random five-fold cross-validation. Specifically, this involved calculating the mean Pearson’s correlation coefficient (PCC) between the predicted and observed values across 50 random tests.

### MVBLUP for predicting SSC trait of Tomato332

The analysis was initiated by assessing the prediction accuracy of MVBLUP for the fruit soluble solid content trait (SSC) of Tomato332. Three distinct views, single nucleotide polymorphisms (SNP), insertions and deletions (InDel) and structural variants (SV), were selected to construct a multi-view prediction.

To highlight the distinct information conveyed by different views, heatmaps of their respective kinship matrices were presented in Fig. 2A. These heatmaps revealed discernible differences in the similarity patterns among the three views’ data. For each view, LASSO, BLUP, and XGBoost were employed independently to predict the SSC trait.

**Fig. 2.**
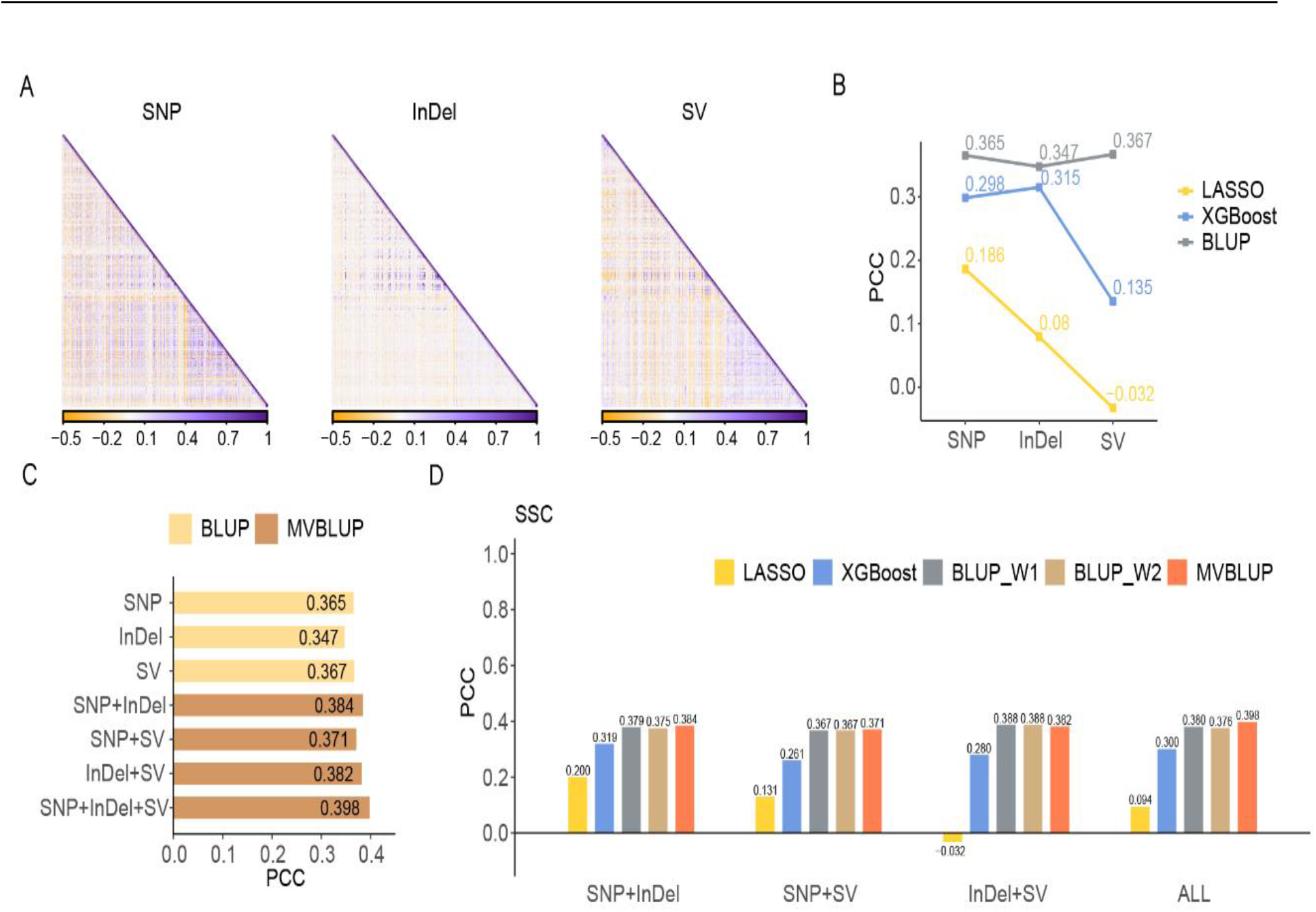
Systematic test results of MVBLUP on the Tomato332 dataset. Heatmaps of the genetic relationships based on SNP, InDel, and SV (**A**). Prediction accuracy of LASSO, XGBoost, and BLUP using single-view data (**B**). Prediction accuracy of BLUP using single-view data and MVBLUP integrating two or three views data (**C**). Prediction accuracy of five methods using three views data (**D**). BLUP_W1: integrating multi-view data with uniform weights and utilizing BLUP for phenotypic prediction; BLUP_W2: integrating multi-view data with weights determined by the average accuracy of five-fold cross-validation on single-view data training sets and employing BLUP for phenotypic prediction.

The results demonstrated that prediction accuracy varied depending on the view data utilized, even when the same method was applied. This further illustrated the distinctiveness of the information carried by different view data (Fig. 2B). Notably, the BLUP method exhibited the highest prediction accuracy across all three types of view data. Given BLUP’s robust performance (Xu et al., 2024), multi-view information was integrated based on BLUP, leading to the development of the MVBLUP method.

Comparisons of BLUP using single-view data, MVBLUP with pairwise integrative views, and MVBLUP with all three integrative views were given (Fig. 2C). Results revealed that integrating additional views improved prediction accuracies. BLUP with a single InDel feature performed the worst (0.347). When MVBLUP incorporated pairwise-view features, the prediction accuracy rose to 0.384. With all three views integrated, MVBLUP achieved the highest accuracy of 0.398.

Including both SNP and InDel feature in LASSO and XGBoost models (Fig. 2D) led to prediction accuracies of 0.20 and 0.319 respectively, making improvements over single-view data. This trend aligns with the results observed in the MVBLUP model (Fig. 2C), suggesting that MVBLUP’s enhanced prediction accuracy is partly attributed to the complementarity of multi-view data.

It is crucial to note that directly integrating multi-view information into a model can sometimes reduce prediction accuracy if the information is redundant or incompatible. For instance, when LASSO and XGBoost methods incorporated the SV feature, their prediction accuracy was respectively 0.218 and 0.163 lower than when using SNP feature (Fig. 2B). Furthermore, the inclusion of the SV feature decreased the prediction accuracies of both methods to 0.131 and 0.261 respectively (Fig. 2D). This underscored the importance of extracting complementary information while excluding redundant and incompatible information during multi-view data fusion.

One solution to this challenge is assigning weights to the multi-view data entering the model. Based on BLUP, three weight assignment methods: uniform weights, weights based on the prediction accuracy of the training set, and optimal weights learned by a differential evolution algorithm (MVBLUP), were employed for multi-view integration. Comparative analysis revealed that MVBLUP outperformed the other two methods, achieving a 0.022 improvement in prediction accuracy (Fig. 2D). Consequently, we adopted the MVBLUP approach for integrating multi-view data.

### MVBLUP for predicting four traits of Rice210

For the dataset Rice210, three distinct types of omics data, genomic (G), gene expression (E) and metabolomic (M), were meticulously selected and seamlessly integrated into a MVBLUP framework to predict four traits: grain number per panicle, 1000 grain weight, yield per plant, and tiller number per plant. Notably, MVBLUP demonstrated remarkable prediction accuracy, outperforming the single-view BLUP method for three out of the four traits being assessed (Fig. 3). Specifically, MVBLUP surpassed the single-view BLUP method by approximately 0.05 in predicting grain number per panicle. For the yield per plant trait, MVBLUP achieved an impressive prediction accuracy of 0.725, which has been significantly improved by 0.296 compared to the single-view BLUP using genomic data alone.

**Fig. 3.**
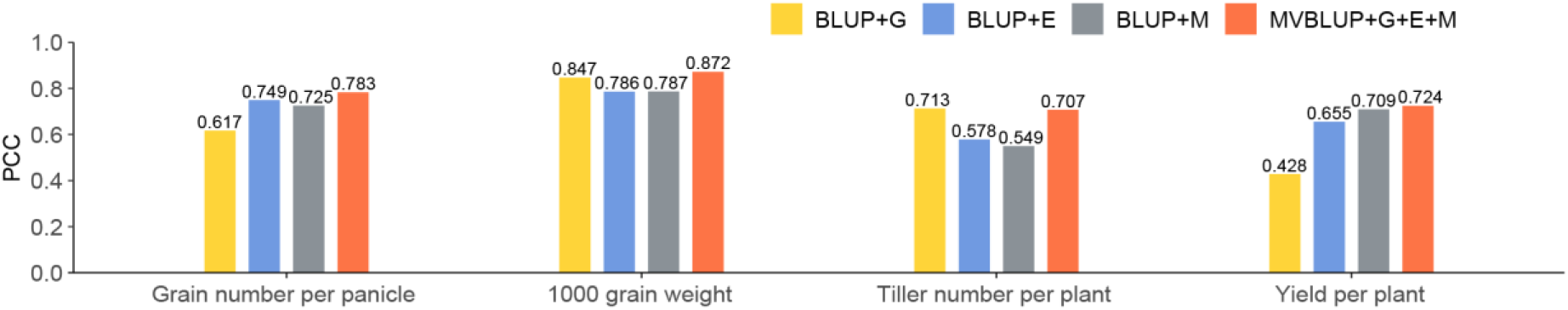
Prediction accuracy of MVBLUP and BLUP using single-view data on the Rice210 dataset. G, genomic data; E, gene expression data; M, metabolomic data.

However, it is worth mentioning that the results for the tiller number per plant trait were somewhat unexpected. In this case, MVBLUP’s accuracy of 0.707 was slightly lower than that of the single-view BLUP with genomic data which was 0.713.

### MVBLUP for predicting eight traits of Maize368

In the context of the Maize368 dataset, MVBLUP was employed to predict eight diverse traits: cob diameter, ear diameter, ear leaf width, ear length, ear leaf length, heading date, pollen shedding and silking time. This prediction was facilitated by the integration of three-view data, comprising genomic (G), gene expression (E) and metabolomic (M). MVBLUP demonstrated remarkable prediction accuracies for seven of these traits (Fig. 4). Specifically, for the trait of heading date, MVBLUP surpassed the peak prediction accuracy of the single-view BLUP method by a margin of approximately 0.041. In the case of silking time, MVBLUP achieved an accuracy of 0.632, marking a 0.073 increase in accuracy compared to the single-view BLUP utilizing genomic data. The only trait where MVBLUP’s performance was somewhat less impressive was ear leaf length, with a prediction accuracy of 0.609, which was marginally lower than the accuracy achieved by the single-view BLUP with genomic data.

**Fig. 4.**
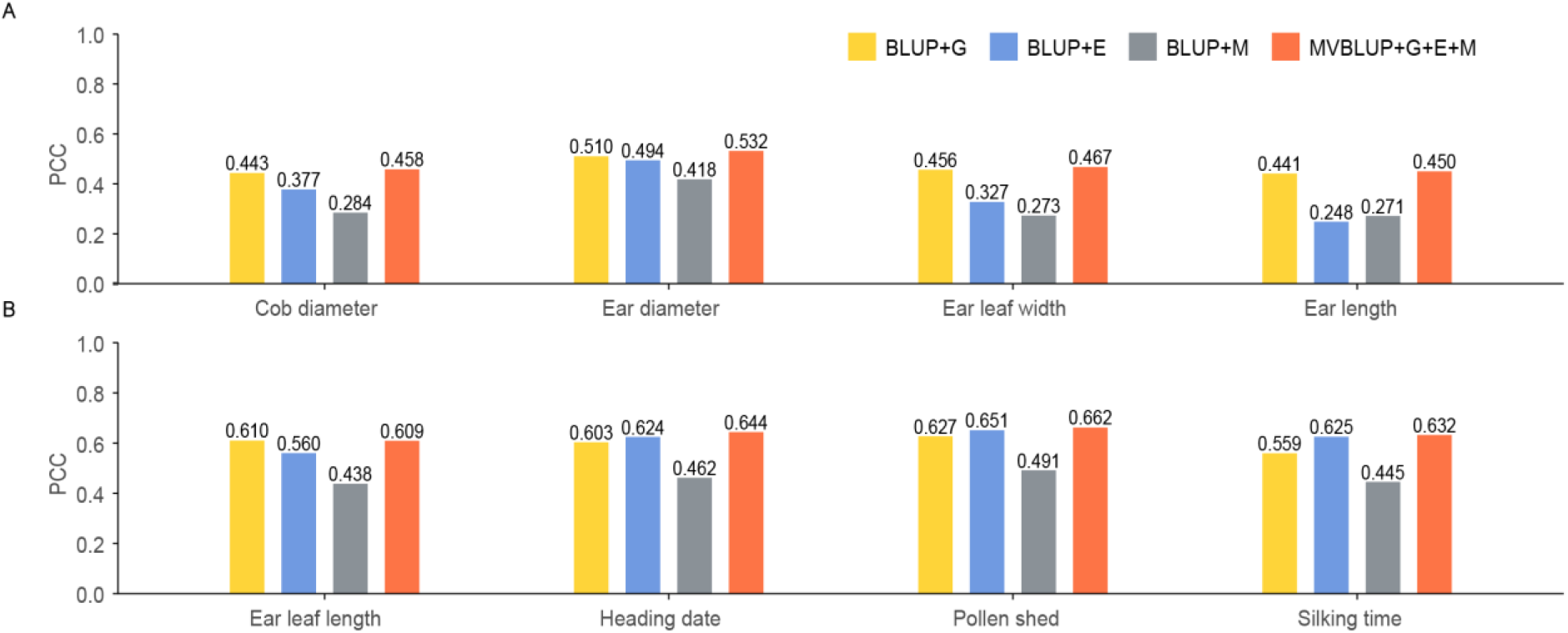
Prediction accuracy of MVBLUP and BLUP using single-view data on the Maize368 dataset. G, genomic data; E, gene expression data; M, metabolomic data.

### MVBLUP for predicting six traits of Maize282

For the Maize282 dataset, MVBLUP was utilized to predict six traits: plant height, weight of 20 kernels, node number below ears, days to anthesis, ear height and leaf width. This prediction was enabled by the integration of eight views, including genomic data (G) alongside gene expression data sourced from seven distinct tissues (E1-E7). MVBLUP exhibited superior prediction accuracy for five out of the six traits when compared to BLUP utilizing individual views (Fig. 5). Notably, MVBLUP surpassed the peak prediction accuracy of the single-view BLUP method for the trait of node number below ears by a margin of approximately 0.039. For the trait of weight of 20 kernels, while MVBLUP’s accuracy of 0.494 was slightly lower than the optimal accuracy of 0.506 achieved by BLUP with the E3 view (gene expression data from the tissue of the third leaf from the base), it still exceeded the performance of the single-view BLUP method, which utilized genomic data, by a notable margin of approximately 0.026.

**Fig. 5.**
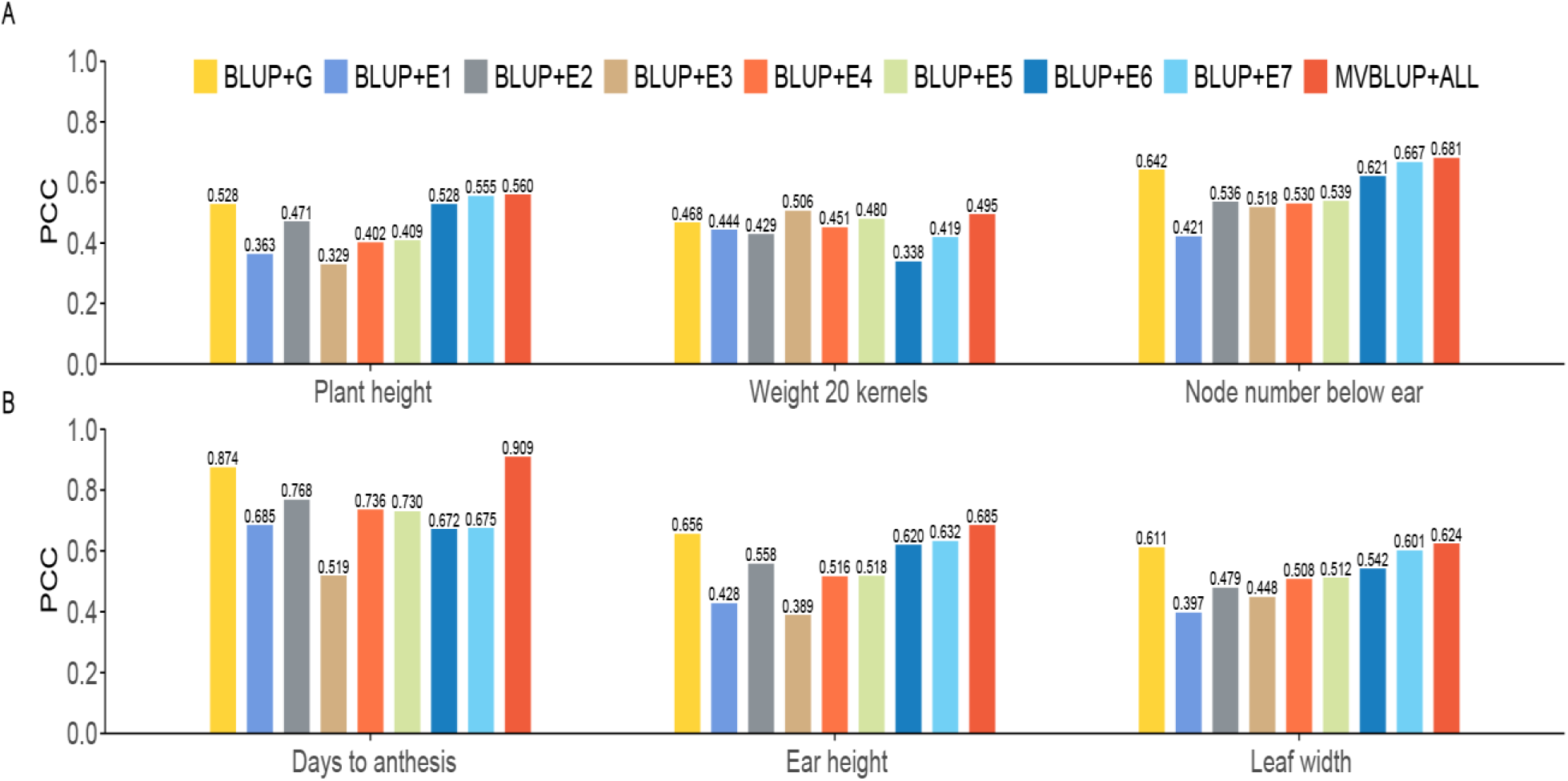
Prediction accuracy of MVBLUP and BLUP using single-view data on the Maize282 dataset. G, genomic data; E1-E7, gene expression data from seven tissues respectively, including germinating root, germinating shoot, third leaf from the base, third leaf from the top, adult leaf collected during the day, adult leaf collected at night and mature kernel.

The efficiency of MVBLUP in the Maize282 dataset was determined by the optimal weight calculated by the DE algorithm. To demonstrate the convergence of the DE algorithm, the iteration process of MVBLUP on Maize282 was shown (Fig. 6). For this dataset, the initial population size was set as 40, with eight weight parameters needing optimization. Fig. 6 revealed that the algorithm converged steadily to the optimal solution as the iteration progressed. Although the maximum iteration number of the DE algorithm was set at 50, the algorithm stabilized after 43 iterations. The early stopping mechanism, which was triggered when the maximum error of the cost function value at two adjacent iteration points fell below a specified tolerance, was instrumental in saving computational cost.

**Fig. 6.**
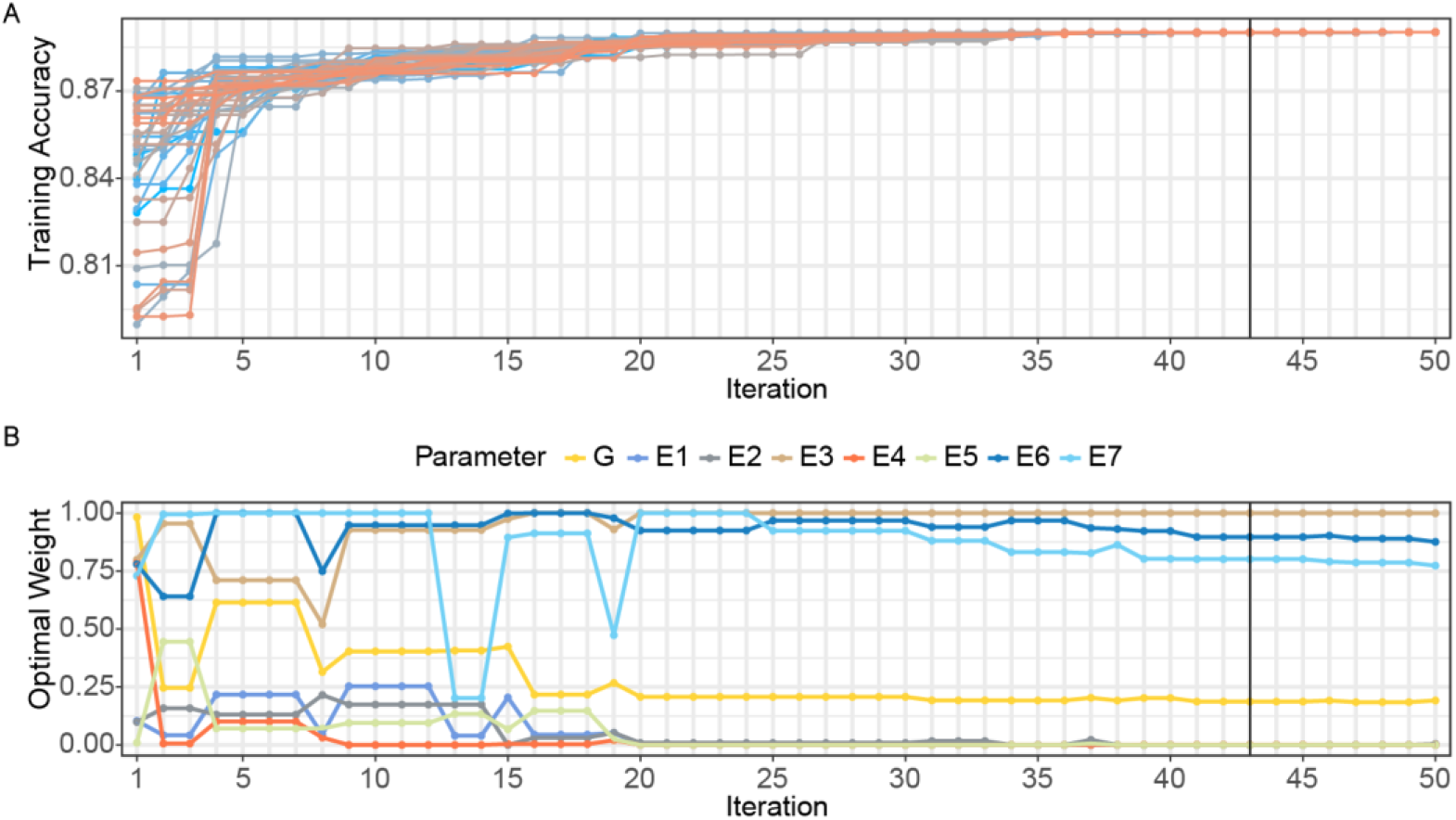
Changes in prediction accuracy and weights during the learning process of the MVBLUP algorithm. The training process of integrating eight single-view data sets G, E1, E2, E3, E4, E5, E6, E7 from Maize282 using the MVBLUP method, exemplified by the maize flowering days phenotype (**A**). The changes in optimal weights of the eight views data during the training process (**B**). G, genomic data; E1-E7, gene expression data from seven tissues respectively, including germinating root, germinating shoot, third leaf from the base, third leaf from the top, adult leaf collected during the day, adult leaf collected at night and mature kernel.

To further validate the effectiveness of the MVBLUP method, numerical experiments analogous to those performed on the Tomato332 dataset were conducted using the Rice210, Maize368, and Maize282 datasets. These experiments involved comparisons with LASSO, XGBoost, BLUP integrating each view with uniform weights (BLUP_W1), and BLUP integrating each view with fixed weights determined by the average accuracy of five-fold cross-validation on single-view data training sets (BLUP_W2). The results of these comparisons were presented in Fig. S1, Fig. S2 and Fig. S3, which showed that MVBLUP outperformed the other methods in most cases. Additionally, the average running time of MVBLUP was significantly faster than the XGBoost method on the Rice210, Maize368, and Maize282 datasets, with average running times of 28.3 seconds, 68.9 seconds, and 283.8 seconds, respectively (Table S1). However, it is worth noting that MVBLUP required the longest average running time on the Tomato332 dataset, at 58.5 seconds.

## Discussion

In this study, we aimed to enhance phenotype prediction by integrating multiple data views through the application of the MVBLUP method. Compared with existing multi-view integrating methods that assign equal importance to each view, MVBLUP offers greater interpretability by an adaptively adjusting weights strategy adjusted with the DE algorithm to quantify the contribution of different views. The strengths of MVBLUP lie in the complementary information derived from multi-view data, the accuracy of single-view models, and the synergistic integration of various views via weights learned by the DE algorithm. These advantages may vary across populations, traits, and datasets.

Recently, a GA-BLUP method, which combines BLUP with a genetic algorithm (GA) to select trait-related markers, has emerged as a highly precise genomic prediction method, particularly for traits with low heritability (Xu et al., 2024). MVBLUP shares some similarities with GA-GBLUP in that both utilize evolutionary algorithms for selection. Specifically, MVBLUP employs DE for selecting multi-view data, whereas GA-GBLUP uses GA for marker selection. Although DE and GA share fundamental operators like mutation, crossover and selection, they differ in key aspects. Notably, the mutation operator in DE fundamentally differs from that in GA. In GA, mutation typically involves randomly altering individual bits or genes in a chromosome, whereas DE employs a differential mutation strategy that explores the search space more efficiently and effectively. In addition, GA excels in discrete optimization problems due to its encoding and decoding mechanism tailored for combinatorial optimization, whereas DE performs better in continuous optimization problems, which are more suitable for integrating multi-view data in this study.

MVBLUP demonstrated superior prediction accuracy compared to BLUP, LASSO and XGBoost, both with single-view and multi-view data, across four datasets, highlighting its effectiveness and applicability. However, it is worth noting that MVBLUP slightly underperformed in three specific cases. For instance, in predicting the tiller number per plant trait of the Rice210 dataset, MVBLUP’s accuracy (0.707) was marginally lower than that of single-view BLUP with genomic data (0.713). We hypothesize that the slight decrement in performance for certain traits could be attributed to discrepancies in data distributions between the training and test datasets, which may lead to the model excelling on the training set but struggling with generalization to the test set. During MVBLUP’s learning process, optimal weights were selected based on the highest fitness function value on the training set. However, in some instances, despite achieving optimized training performance, the testing accuracy fell short of expectations. To illustrate this phenomenon, we randomly divided the Rice210 dataset into training and test sets 50 times and calculated the accuracy using MVBLUP and single-view BLUP based on genomic data after each division. MVBLUP exhibited better training accuracy than single-view BLUP method in each training set (Fig. S4A). However, in terms of test performance, MVBLUP showed lower prediction accuracy in some test sets (Fig. S4B). This comparison underscores the potential discrepancy between training performance and actual testing outcomes, which could be partially mitigated by increasing the dataset’s sample size.

Moreover, a deeper exploration of MVBLUP’s enhancements is crucial for future research. In this study, we assigned weights using the DE algorithm, commonly used in vast and complex parameter spaces. However, with a maximum of eight multi-view data sources, the full advantages of DE were not fully demonstrated. Importantly, our MVBLUP framework is inherently scalable to accommodate a broader range of multi-view scenarios. In breeding, MVBLUP offers a solution for efficiently integrating multi-view data including genomic data, Red Green Blue (RGB) images, and spectrum image-based phenomic data. The use of Unmanned Aerial Vehicles (UAVs) for high-throughput phenotyping will drastically accelerate the collection of multi-view data related to plant physiological status throughout the growth period, achieving both efficiency and cost-effectiveness. Alternatively, if focus solely on genomic data, we can categorize the data by chromosomes and assign weights accordingly. Furthermore, we can assign weights to each SNP marker and employ the DE method to learn and optimize these weights. As an evolutionary algorithm, DE has the potential to avoid local optima but may compromise computational efficiency. Therefore, future research could explore efficient alternatives, such as improved DE strategies with accelerated convergence (Bilal et al., 2020).

## Materials and Method

### Data sources

The Tomato332 dataset comprises 332 materials from three tomato subspecies: currant tomato (PIM), cherry tomato (CER), and large-fruited cultivated tomato (BIG). The genotype data for this dataset is a call set named TGG1.1-332 from the tomato graph pangenome, which encompasses 6,971,059 SNPs, 657,549 InDels, and 54,838 SVs. A crucial phenotypic trait in this dataset is the fruit soluble solids content (SSC), which is significant for both yield and flavor (Zhou et al., 2022). By applying Principal Component Analysis (PCA) to the genotype data, the dataset was reduced to 220 SNPs, 289 InDels, and 277 SVs, which were then used as a multi-view data for predicting the SSC trait (Wang et al., 2023).

The Rice210 dataset consists of 210 recombinant inbred lines (RILs), obtained through the crossing two rice varieties Zhenshan 97 and Minghui 63 (Hua et al., 2003). Sequencing of these RILs resulted in the identification of 270,820 high-quality SNPs and 1,619 bins based on recombination breakpoints, serving as the genotype data (Yu et al., 2011). Ribonucleic Acid (RNA) was extracted from the flag leaves of the RILs during the heading stage between 8:00 and 9:30 AM, and the expression levels of 24,994 genes were quantified using a microarray sequencing platform, providing transcriptomic data (Wang et al., 2014). Metabolomic data included 1,000 metabolites sourced from two tissues: flag leaves at heading stage and seeds 72 hours post-germination (Gong et al., 2013). Four key agronomic traits—yield per plant (Yield), tiller number per plant (Tiller), grain number per panicle (Grain), and 1,000-grain weight (KGW)—were collected in 2008 and 2009 from a field experiment conducted at the Farm of Huazhong Agricultural University in Wuhan, China (Yu et al., 2011).

The Maize368 dataset includes 368 maize inbred lines derived from a broadly representative association mapping population encompassing tropical, subtropical, and temperate germplasm (Yang et al., 2011). These inbred lines were genotyped using four different genotyping platforms, resulting in the identification of 1.25 million high-quality SNP markers as the genotype data (Liu et al., 2017). RNA was extracted from immature kernels at 15 days post-pollination and sequenced to obtain expression levels for 28,769 genes (Fu et al., 2013). Additionally, metabolic profiling of mature maize kernels led to the identification of 749 non-targeted metabolites (Wen et al., 2014). This study utilized eight agronomic traits previously analyzed in a Genome-Wide Association Studies (GWAS) study (Yang et al., 2014), including cob diameter, ear diameter, ear leaf width, ear length, ear leaf length, heading date, pollen shed and silking time. These traits were collected across five environments, and their average values were used for phenotypic prediction (Yang et al., 2014).

The Maize282 dataset comprises 282 maize inbred lines sourced from a US maize association mapping panel (Flint-Garcia et al., 2005). Genotyping of these inbred lines was conducted using the Illumina MaizeSNP50 BeadChip, resulting in the identification of 50,878 high-quality SNP markers as the genotype data (Ganal et al., 2011). RNA extraction and sequencing were performed on seven different tissues at specified times and locations, including the base and tip of the third leaf collected between 10:30 and 12:00, the root of a 2 cm germinated seedling and the entire shoot of the germinated seedling collected between 11:00 and 13:00 on the day of germination, developing kernels post-pollination collected between 11:00 and 13:00, and mature leaf samples collected from a 1 cm section adjacent to the midrib of the second leaf below the tassel between 11:00 and 13:00, and also between 23:00 and 1:00 (Kremling et al., 2018). Quantitative analysis of messenger RNA (mRNA) expression levels provided transcriptomic data. Six important agronomic traits, including plant height, weight of 20 kernels, node number below ear, days to anthesis, ear height and leaf width, were used in this study (Flint-Garcia et al., 2005).

## Method

MVBLUP is a prediction model that builds upon the BLUP framework and incorporates the DE algorithm to dynamically determine the optimal integrating weight of multi-view features. To ensure comprehensiveness, we first introduce the principle of BLUP and DE.

### Best linear unbiased prediction

The BLUP approach relies on a mixed linear model. The fundamental model could be articulated as:

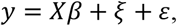

where:

- *Y* is an *n* × 1 vector of phenotypic values of a quantitative trait for *n* individuals.
- *X* is an *n* × *p* design matrix.
- *β* is a *p* × 1 vector of fixed effects.
- *ξ*∼*N*(0, *Kσ*^2^) is a *n* × 1 vector of random effects.
- 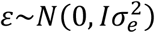) is a *n* × 1 vector of residual errors.
- *K* represents the relationship between individuals.
- *σ*^2^ and 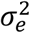 are variance associated with random effects and residual errors, respectively.
- *I* is an identity matrix.

The random effects vector *ξ* can be derived using:

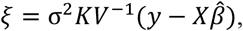

where 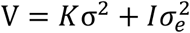 *and* 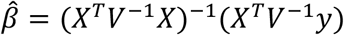.

### The similarity function and kinship matrix

Let *x*_*i*_ denotes the input feature vector of the *i*th individual (for example, the expression level of the *i*th individual). The similarity between the *i*th and the *j*th individual is defined as:

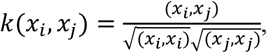

where (*x*_*i*_, *x*_*j*_) represents the inner product of the vector *x*_*i*_ and *x*_*j*_.

Based on the similarity function, we define a single view initial kinship matrix *A*_0_ as: 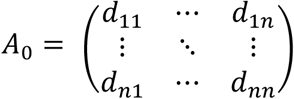, where *d*_*ij*_ = (*x*_*i*_, *x*_*j*_) and *n* is the number of individuals. Further, we can define the single view kinship matrix *A* as: 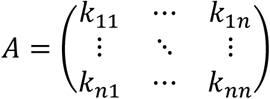 where *k*_*ij*_ = *k*(*x*_*i*_, *x*_*j*_). Finally, we define the relationship matrix *K* which integrates multiple views of data as: 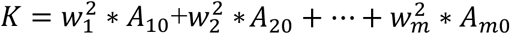, where 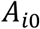 is the *i*th view initial kinship matrix, *m* is the number of view and *w*_*j*_ is the weight of the *j*th view data. Here the new multi-view relationship matrix can be seen as a generalization of single view kinship matrix.

### Differential evolution algorithm

In this study, DE is employed to identify the optimal weights for each view’s initial kinship matrix. The DE process encompasses initialization, mutation, crossover, and selection. The steps are detailed as follows:

#### Step 1. Initialization

An initial population 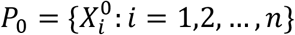 is generated as follows:

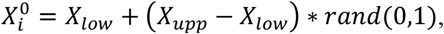

where *n* is the size of population, and *X*_*low*_ and *X*_*upp*_ are lower and upper bounds of search space, respectively.

#### Step 2. Mutation

The *i*th mutant individual in the *g*th vector generation is created according to the following:

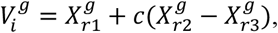

where 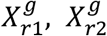 and 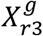 are the randomly selected individuals from the parent population, *r*1, *r*2,*r*3 ∈ {1,2, …, *n*}, and *c* ∈ (0,1) is the scaling factor.

#### Step 3. Crossover

The *i*th crossover individual in the *g*th trial vector is generated as follows,

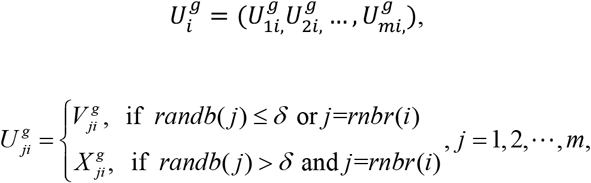

where *randb*(*j*) ∈[0,1] is a uniform random number, *δ* ∈[0,1] is a predefined crossover parameter, *ranbr*(*i*) ∈{1, 2,, *m*} is an index selected randomly.

#### Step 4. Selection

Determine whether the trial vector in the crossover step should be included in next generation by a strategy as follows,

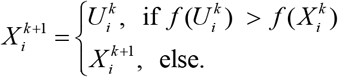

Where *f* (*X*) is a fitness function.

#### Step 5. Stopping criterion

If maximum error of the fitness function value between 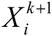 and 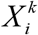 is less than tolerance *ε* (i.e., an early stopping mechanism), or the maximum iteration count *K* is reached, then DE algorithm will be stopped.

Prior to training, five parameters must be set: population size *n*, scaling factor *c*, crossover parameter *δ*, tolerance *ε* and maximum iteration parameter *K*. In our experiment, Population size *n* is set as five times number of feature views. Specifically, *n* = 15 for Tomato332, Rice210 and Maize368, *n* = 40 for Maize282. Both scaling factor *c* and crossover parameter *δ* are set as 0.5, tolerance *ε* = 0.0001 and maximum iteration number *K* = 50.

### The multi-view best linear unbiased prediction procedure

MVBLUP is a flexible machine learning algorithm that integrates adaptively multi-view data for phenotype prediction. The MVBLUP process involves:

#### Step 1 Normalization

Input vectors are normalized using the Z-score method, and initial weights are assigned randomly to establish the initial population 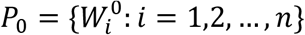 for DE.

#### Step 2 Training. Repeat the following steps *K* times or stop with an early stopping mechanism, Step 2.1

Set *k* = 1. By utilizing the “Mutation and Crossover” steps within the Differential Evolution (DE) algorithm, the *n* weight vectors are renewed. Subsequently, these updated weight vectors are employed to compute *n* multi-view kinship matrices through the application of the similarity function.

##### Step 2.2

Using “Selection” steps in DE algorithm to renew the weight vector, we obtain the initial population of the next generation. The “Selection” step utilizes the average prediction accuracy derived from five-fold cross-validation on the training set as the fitness function, which can be mathematically represented as 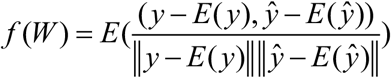, where *E* represents expectation, *y* is the observed value vector for a particular trait, *ŷ* is the predicted value vector for the corresponding trait, and ‖ ·‖ represents the 2-norm, 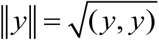.

##### Step 2.3 (Stopping criterion)

If the difference between *f* (*W* ^*k*+1^) and *f* (*W* ^*k*^) is less than tolerance, then optimal weight vector is found, training will be stopped; Else, *k* = *k* + 1, go to Step 2.1.

#### Step 3 (Prediction)

Phenotypes are predicted using the optimized multi view kinship matrix

### Methods used for comparisons

The predictive capabilities of MVBLUP was initially compared with BLUP when utilizing single-view data. Subsequently, two additional prevalent techniques: least absolute shrinkage and selection operator (LASSO) and extreme gradient boosting (XGBoost), were applied to both single view and multi-view datasets for comparative purposes.

LASSO aims to identify an optimal linear model represented as *y*= *X*α + *ε*, where *X* is an *n* × *d* matrix, *y*is a *n* − dimensional vector, *ε* denotes noise and α serves as the weight vector. To determine an appropriate α, LASSO can be formulated as the following optimization problem:

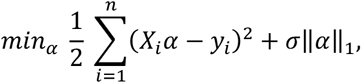

with *σ* being a regularization parameter. In this study, LASSO method was executed using the GLMNET/R package (Friedman et al., 2010).

XGBoost method, on the other hand, focuses on constructing an ensemble of trees, utilizing *K* additive functions to predict the output,

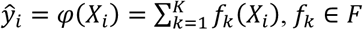

where *X*_*i*_ ∈ *R*^*d*^, *y*_*i*_ ∈ *R, F* = {*f*(*x*) = *ω*_*q*(*x*)_}(*q:R*^*d*^ → *T, ω* ∈ *R*^*T*^) represents the space of regression trees (also called as CART), *q* represents the structure of each tree, *T* is the number of leaves in the tree, each *f*_*k*_ corresponds to an independent tree structure *q* and leaf weight *ω*.

To learn the functions in the model, XGBoost can be framed as the following optimization problem,

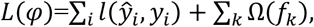

where *l* is a differentiable convex loss function measuring the discrepancy between the prediction 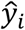 and the target 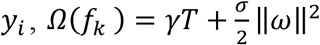 serves as a regularization term penalizing model complexity, γ and *σ* are regularization parameters (Chen and Guestrin, 2016). In this study, XGBoost was implemented using the xgboost/R package.

## Data availability

The MVBLUP algorithm’s demo data, R scripts, and tutorial are available for public access on GitHub at the following link: https://github.com/bjwu555/MVBLUP.

## CrediT authorship contribution statement

**Jianbing Yan, Wenyu Yang** and **Yingjie Xiao**, designed and supervised this study. Bingjie Wu established the MVBLUP. **Bingjie Wu** and **Huijuan Xiong** performed data analysis and wrote the draft. **Lin Zhuo** participated in the evaluation of MVBLUP. **Bingjie Wu, Huijuan Xiong, Lin Zhuo, Wenyu Yang** and **Yingjie Xiao** prepared the manuscript. The authors read and approved the final manuscript.

## Acknowledgments

This work was supported by National Natural Science Foundation of China (32122066, 32201855), STI2030—Major Projects (2023ZD04076).

