## Supplemental figure s1-s4 and supplmental table 1 for "Multi-view BLUP: a promising solution for post-omics data integrative prediction"

**Supplementary figures:**

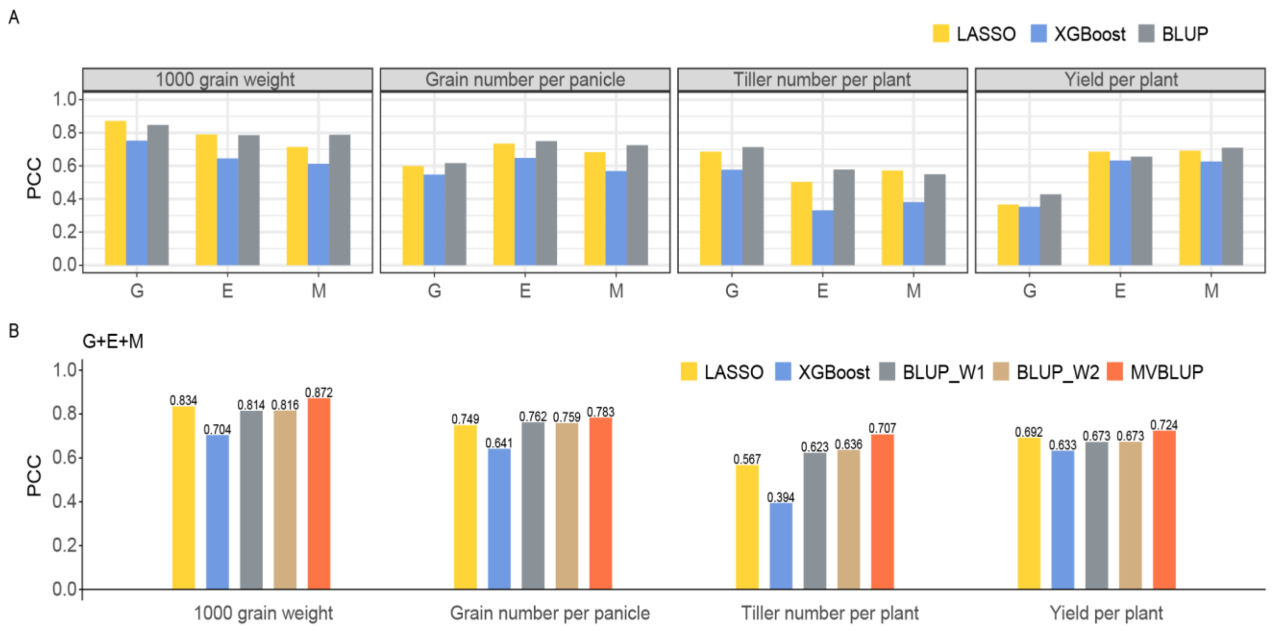


**Fig. S1. Prediction accuracy of single-view and multi-view methods on the Rice210 dataset.** Prediction accuracy of three methods using single-view data (**A**). Prediction accuracy of five methods using three views data for four traits (**B**). G, genomic data; E, gene expression data; M, metabolomic data; BLUP_W1: integrating multi-view data with uniform weights and utilizing BLUP for phenotypic prediction; BLUP_W2: integrating multi-view data with weights determined by the average accuracy of five-fold cross-validation on single-view data training sets, and employing BLUP for phenotypic prediction.

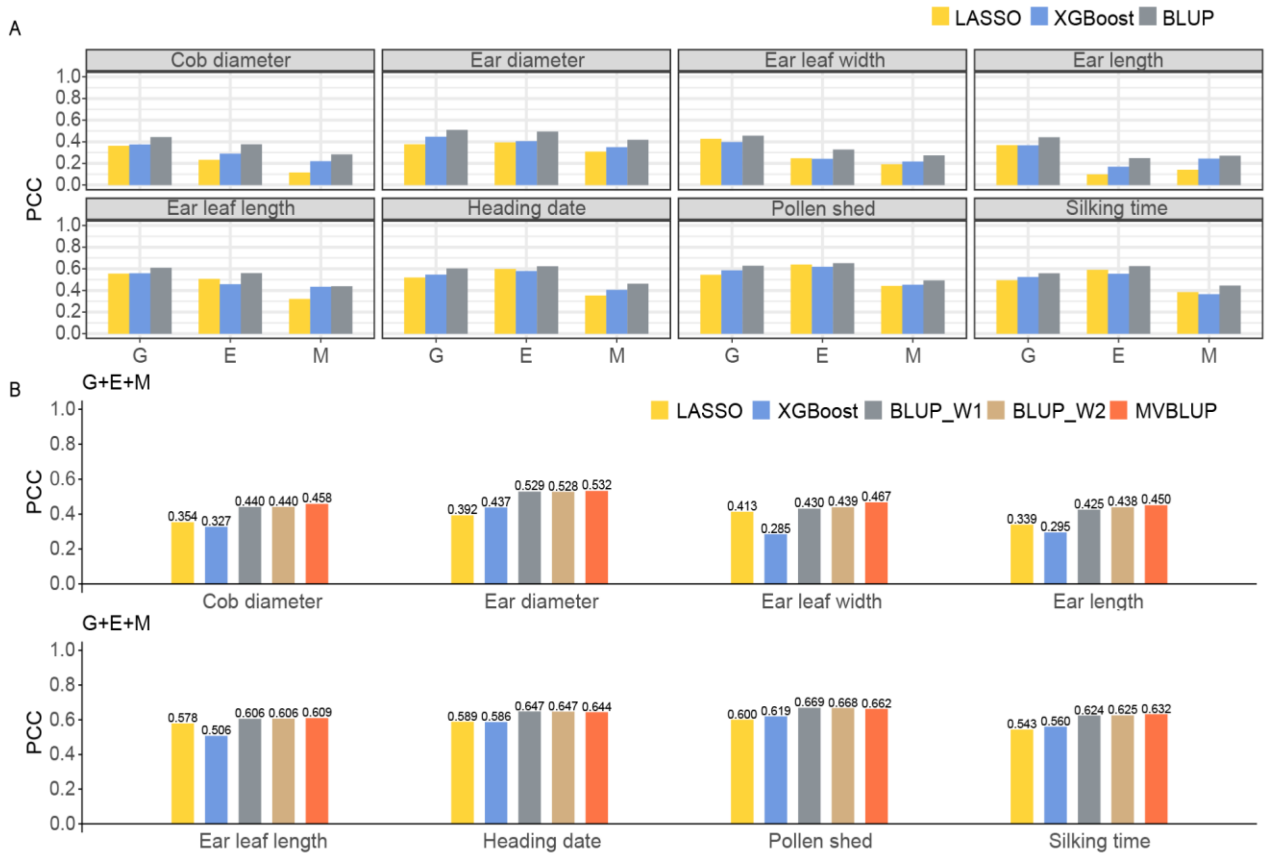


**Fig. S2. Prediction accuracy of single-view and multi-view methods on the Maize368 dataset.** Prediction accuracy of three methods using single-view data (**A**). Prediction accuracy of five methods using three views data for eight traits (**B**). G, genomic data; E, gene expression data; M, metabolomic data; BLUP_W1: integrating multi-view data with uniform weights and utilizing BLUP for phenotypic prediction; BLUP_W2: integrating multi-view data with weights determined by the average accuracy of five-fold cross-validation on single-view data training sets, and employing BLUP for phenotypic prediction.

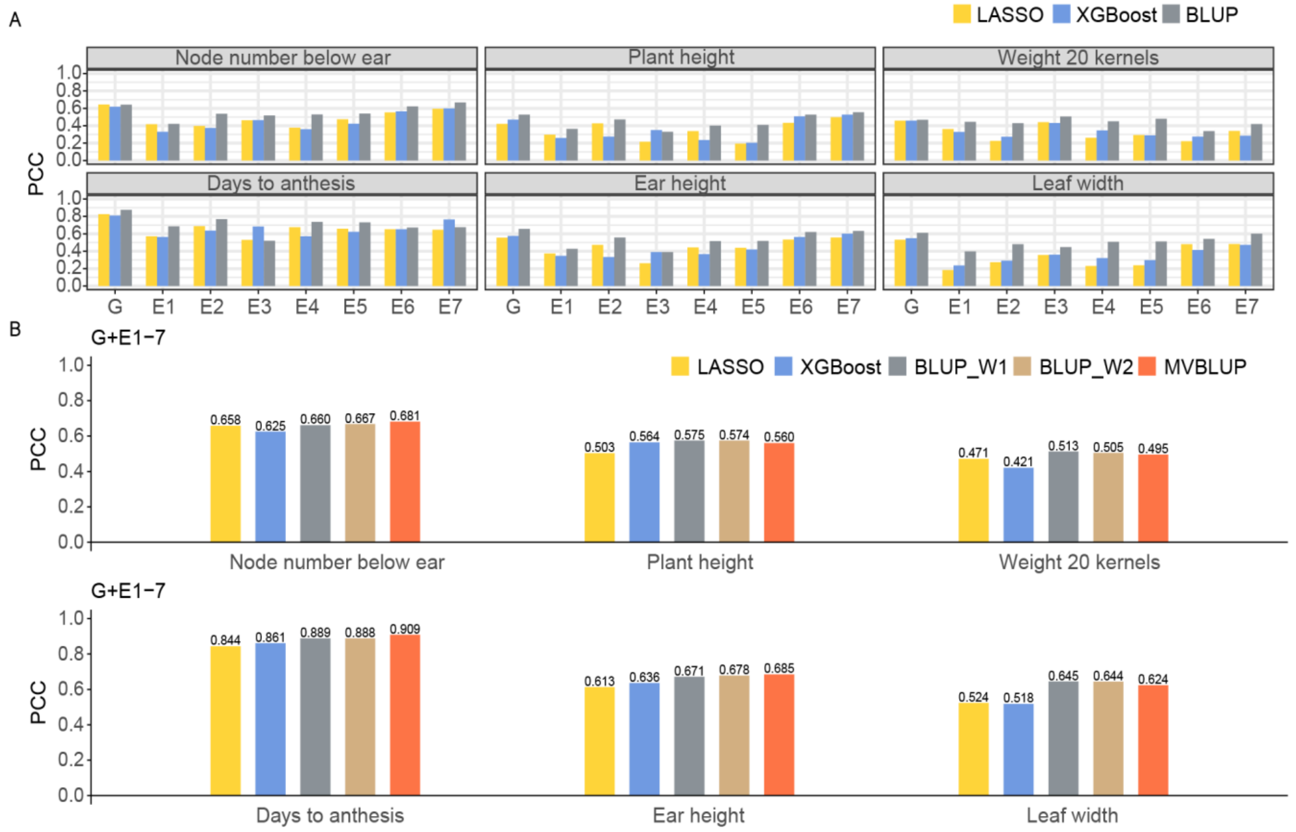


**Fig. S3. Prediction accuracy of single-view and multi-view methods on the Maize282 dataset.** Prediction accuracy of three methods using single-view data (**A**). Prediction accuracy of five methods using three views data for six traits (**B**). G, genomic data; E1-E7, gene expression data from seven tissues respectively, including germinating root, germinating shoot, third leaf from the base, third leaf from the top, adult leaf collected during the day, adult leaf collected at night and mature kernel; BLUP_W1: integrating multi-view data with uniform weights and utilizing BLUP for phenotypic prediction; BLUP_W2: integrating multi-view data with weights determined by the average accuracy of five-fold cross-validation on single-view data training sets, and employing BLUP for phenotypic prediction.

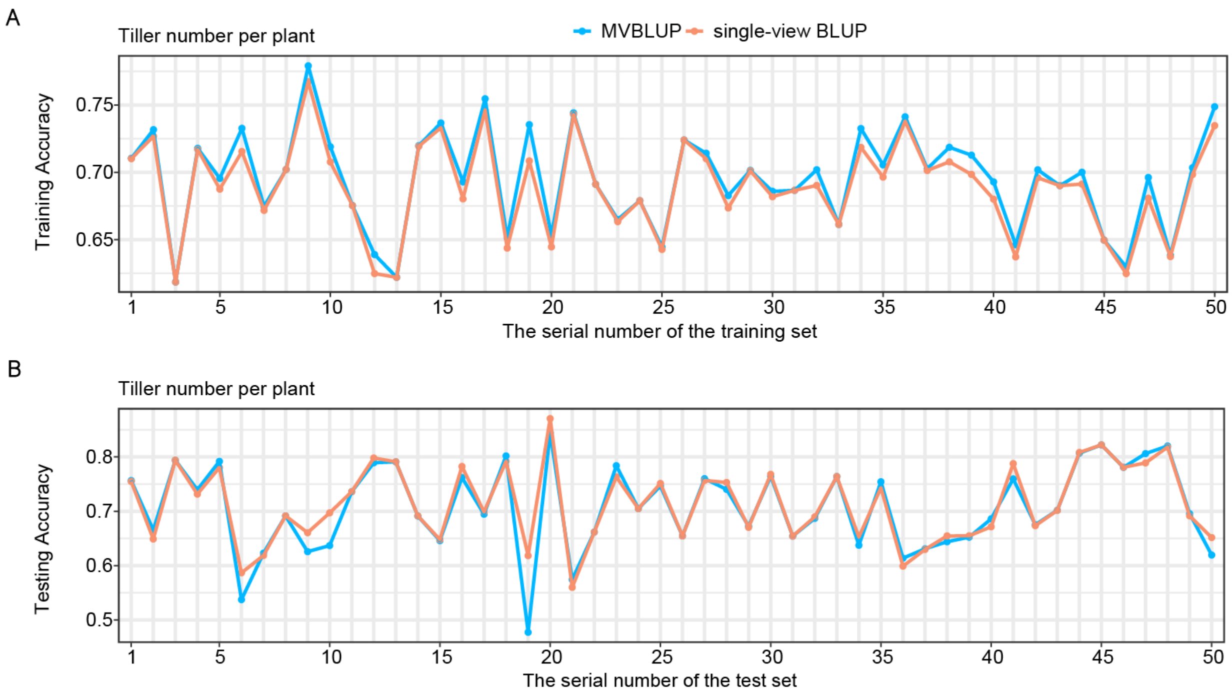


**Fig. S4. Comparison of predictive accuracies for the trait "Tiller number per plant" in the Rice210 dataset using two methods: MVBLUP and single-view BLUP, with the latter incorporating genomic data.** Prediction accuracy of MVBLUP and single-view BLUP across 50 random repetitions on training set (**A**). MVBLUP and single-view BLUP across 50 random repetitions on test set (**B**).

**Supplementary table：**

**Table 1. Comparison of average computational time (sec.)**

| | LASSO | XGBoost | BLUP_W1 | BLUP_W2 | MVBLUP |
|---|---|---|---|---|---|
| Tomato332 | 0.1631 | 10.3750 | 0.0435 | 0.4985 | 58.5136 |
| Rice210 | 2.1664 | 153.6318 | 0.0992 | 0.3068 | 28.2721 |
| Maize368 | 15.7274 | 526.8456 | 0.0950 | 0.6287 | 68.9041 |
| Maize282 | 12.4642 | 644.5843 | 0.3396 | 1.1737 | 283.7643 |

BLUP_W1: integrating multi-view data with uniform weights and utilizing BLUP for phenotypic prediction; BLUP_W2: integrating multi-view data with weights determined by the average accuracy of five-fold cross-validation on single-view data training sets and employing BLUP for phenotypic prediction.
